# MBD2/3 lost its methyl-CpG binding ability in multiple families of Holometabola

**DOI:** 10.1101/2025.04.22.650097

**Authors:** Elisa Israel, Zoe Marie Länger, Jacqueline Heckenhauer, Joachim Kurtz, Sonja Prohaska

**Affiliations:** Computational EvoDevo Group, Institute of Computer Science, Leipzig University, Leipzig, Germany; Animal Evolutionary Ecology Group, Institute for Evolution & Biodiversity, University of Münster, Münster, Germany; Department of Terrestrial Zoology, Entomology III, Senckenberg Research Institute and Natural History Museum Frankfurt, Frankfurt, Germany; Joint Institute for Individualisation in a Changing Environment, University of Münster and Bielefeld University, Münster and Bielefeld, Germany

**Keywords:** methyl-CpG binding domain, insects, CpG methylation, isoforms

## Abstract

DNA methylation is sparse in insects, compared to vertebrates. This reduction is even more pronounced in Holometabola, where the loss of DNA methyltransferases DNMT1 and DNMT3 has occurred several times. In some Holometabola, DNA methylation is lost entirely. Methyl-CpG-binding domain (MBD) proteins bind methylated CpGs and are therefore important readers of these epigenetic marks. We hypothesize that the evolutionary reduction of genome-wide methylation may be paralleled by changes to MBD proteins. Among insects, only a single MBD family member, MBD2/3, is known. Two isoforms of MBD2/3 have been identified in *Bombyx mori*, MBD2/3-L and MBD2/3-S. The long isoform MBD2/3-L contains a complete MBD domain spanning the first two exons, whereas the short isoform MBD2/3-S lacks the second exon and therefore half of its MBD domain. It has been reported that only the MBD2/3-L isoform is able to bind methyl-CpGs.

In this study, we analyzed transcriptomic and genomic sequence data across holometabolous orders to identify MBD2/3 genes and their isoforms. Our findings reveal that MBD2/3 is highly conserved in sequence and gene structure. While both isoforms are present in most hemimetabolous orders, the long isoform MBD2/3-L, capable of binding methylated CpG, has been lost in multiple holometabolous orders. The results suggest that MBD2/3 has lost its ability to bind methyl-CpG in several insect orders, with several independent losses in Holometabola. This occurred through different changes to the MBD domain, from sequence divergence within the domain to the absence of half of the MBD domain. These losses may be linked to the reduced levels of CpG DNA methylation.

## Introduction

DNA methylation is a widespread epigenetic modification found across most clades in the tree of life (Nasrullah et al. 2022). Despite its ancient evolutionary origins, a drastic disparity in the distribution and prevalence of DNA methylation is observed between vertebrates and invertebrates (Keller, Han, and Yi 2016).

From their last common ancestor, the CpG methylation patterns changed dramatically between invertebrates and vertebrates. From an ancestral state of likely low levels of intragenic CpG methylation (de Mendoza et al. 2019), vertebrate genomes show a global expansion of methylation patterns, including methylation at transposable elements (TEs) and regulatory regions, where methylation outside the gene body is linked to gene silencing and transcriptional control (repression) (Zemach et al. 2010). In contrast, in invertebrates, DNA methylation is sparse and predominantly enriched in gene bodies, mainly exons of actively expressed genes, where it likely plays a role in gene expression stabilization (Dixon and Matz 2022). DNA methylation levels drop further at the origin of holometabolous insects (Provataris et al. 2018). In some species or even whole orders, CpG methylation has been lost entirely. Prominent examples include model organisms such as *Tribolium castaneum* (Coleoptera) (Schulz et al. 2018) and the order of Diptera (Bewick et al. 2017), which exhibit no functional CpG methylation. Despite the differences in location and prevalence of CpG methylation, the general CpG methylation toolkit is evolutionarily conserved and consists of writers, readers and erasers of methylation marks (Rausch, Hastert, and Cardoso 2020). DNA methyltransferases (DNMTs) either methylate previously unmethylated CpGs *de novo* (DNMT3) or maintain methylation patterns during DNA replication and repair (DNMT1) (Lyko 2018). Notably, some invertebrates have lost either DNMT3 or both, DNMT1 and DNMT3. However, the loss of DNMT3 does not necessarily coincide with the loss of CpG methylation, while species that lost DNMT1 also lost DNA methylation (Engelhardt et al. 2022). The reduction or loss of DNA methylation can be expected to have consequences not only for the presence of writers of DNA methylation, but also for readers of methylation or other interaction partners.

Readers of DNA methylation fall into three main classes: the Kaiso and Kaiso-like protein family with a BTB-POZ domain, proteins containing an SRA (SET and RING finger associated) domain, and the methyl-CpG-binding domain (MBD) protein family (Menafra and Stunnenberg 2014). MBD proteins possess a conserved methyl-CpG binding domain essential for recognizing methylated DNA. Before the vertebrate-invertebrate divergence, the ancestral MBD-containing protein was likely an ancestral MBD2/3 (Hendrich and Tweedie 2003). In invertebrates, MBD2/3 is the sole MBD protein, whereas vertebrate genomes expanded the MBD family through gene duplication events. This gave rise to MBD1-6 and MeCP2, with distinct methylation-dependent and independent functions (Baubec et al. 2013). For instance, mammalian MBD2 and MBD3 share approximately 70% sequence identity but differ in intron size and DNA binding affinity (Hendrich and Tweedie 2003). A point mutation in the MBD domain of vertebrate MBD3 alters its binding preference from methylated CpG sites to unmethylated and hemimethylated sites (Hendrich and Tweedie 2003). The conserved MBD domain itself, around 70 amino acids long, is necessary and sufficient to specifically recognize mCpG (Du et al. 2015). It forms a α/β sandwich structure comprising four β-strands, an α-helix and a hairpin loop. Together with the loop, which links β_2_ and β_3_, the twisted β-sheet is embedded in the major groove and stabilized by α_1_ at the target DNA site. A hydrophobic patch formed by residues (Val-20, Arg-22, Tyr-34, Arg-44, and Ser-45) in β_2_ and β_3_ is critical for the MBD-mCpG interaction (Ohki et al. 2001).

While only MBD2/3 exists in invertebrates, two isoforms of MBD2/3 have been reported in several invertebrate species. In the lepidopteran species *Bombyx mori* and *Spodoptera litura*, both possessing DNMT1 and CpG methylation, a long and a short isoform were identified (Fu et al. 2023). Only the long isoform, MBD2/3-L, was found to be able to bind mCpGs. The short isoform, MBD2/3-S, lacking α_1_ and β-strands downstream of β_2_ and β_3_, was found to be unable to bind mCpGs. This was attributed to the stabilizing effects of the missing structural components, making the MBD loose and disordered (Fu et al. 2023; Xu et al. 2021). Also in *Drosophila*, two isoforms were identified, one of which lacks part of the MBD and the other one specifically binds CpT/A (Marhold et al. 2004).

Most MBD proteins can recruit and interact with histone deacetylases (Le Guezennec et al. 2006), thereby mediating transcriptional repression. Vertebrate MBD2 or MBD3 and likely invertebrate MBD2/3 are key components of the nucleosome remodeling and deacetylase (NuRD - alias NDR or Mi-2) complex, involved in transcriptional repression through the recruitment of histone deacetylase (HDAC1/2) to methylated sites (Le Guezennec et al. 2006). As part of the NuRD complex, MBD2/3 serves as a scaffold by linking CHD 3/4/5 (chromodomain helicase DNA binding domain protein CHD 3/4/5), an ATPase, and GATAD2A/B (alias p66 alpha and beta), a transcription factor. Histone chaperones RbAp46/48 and DNA-binding proteins MTA1/2/3 are also associated with the canonical NuRD complex, while zinc finger proteins SALL1/4 occur in a cell-specific context (Hoffmann and Spengler 2019; Allen, Wade, and Kutateladze 2013). The complex plays essential roles in gene transcriptional regulation, genome stability, chromatin remodeling, DNA repair, cell cycle progression, stem cell differentiation, and cerebral corticogenesis (Sokpor et al. 2018; Hoffmann and Spengler 2019; Lai and Wade 2011). MBD3 deletion leads to early embryonic lethality in mice and stem cells lacking MBD3 fail to differentiate (Kaji et al. 2006).

Apart from recruiting histone deacetylases, as known in mammals, Xu et al. (2021) found that in *B. mori* MBD2/3 recruits the acetyltransferase Tip60, enhancing gene expression rather than repressing it. Mechanistically, MBD2/3 binds mCpG close to the TSS, and recruits the histone acetylase Tip60, which acetylates H3K27. It is important to note that only the insect coiled-coil domain of MBD2/3, but not the vertebrate coiled-coil domain, can interact with Tip60 according to Xu et al. (2021).

Interestingly, invertebrates that lost functional DNA methylation retained MBD2/3. Cramer et al. (2017) demonstrated that MBD2/3 and GATAD2A/B bind each other even in the absence of methylation, as shown in *Drosophila* and sponges. This suggests that the role of MBD2/3 as a component of the NuRD complex may be independent of its ability to bind methylated CpGs, potentially explaining its evolutionary conservation.

In invertebrates, not only in those lacking DNA methylation, MBD2/3 appears to perform methylation-independent functions. In the planarian species *Schmidtea mediterranea*, MBD2/3 lacks conserved residues critical for methylated DNA binding. However, it is involved in adult stem cell pluripotency in certain cell lineages, potentially through its role in the NuRD complex (Jaber-Hijazi et al. 2013). In *D. melanogaster*, NuRD formation is regulated during embryogenesis. Here, MBD2/3-S primarily interacts with NuRD components (Marhold et al. 2004). In *T. castaneum*, MBD2/3 is a single-exon gene and lacks the ability to bind methylated DNA (Song et al. 2020). Still, knockdown is lethal in last instar larvae and knockdown in pupae leads to reduced fecundity and impaired ovaries of emerging adults. Fu et al. (2023) examined MBD2/3-L in 14 insect species, including hemimetabolous and holometabolous insects (e.g., the neuropteran *Chrysoperla carnea* and lepidopteran species) and confirmed the presence of the long isoform (MBD2/3-L). However, it has to be noted that the current data is insufficient to draw definitive conclusions about the completeness or absence of the DNA methylation toolkit in certain insect orders. The repeated loss and variability in the toolkit components within certain orders (e.g., Coleoptera) underscore the need for a comprehensive screening of holometabolous orders to fully understand the evolution of DNA methylation systems. Therefore, we investigated the gene structure and transcript isoforms of *mbd2/3* using publicly available genome and transcriptomic data, respectively. We thoroughly investigate all orders of Holometabola and representatives of all orders of hemimetabolous insects. We thereby expand the previously scattered knowledge about the gene structure and completeness of MBD proteins to examine the potential coevolution of the major reader of methylated CpGs and CpG methylation in a supra-order where methylation is only present at low levels, or was lost entirely.

## Materials and Methods

### Data acquisition and species selection

For the holometabolous orders, which are the focus of our study, publicly available genome and transcriptome data from NCBI genome and the NCBI Transcriptome Shotgun Assembly database (TSA), respectively, were retrieved. With the aim to represent a broad phylogenetic range, one or two species per available family of each taxonomic order, depending on data quality, were selected. This resulted in a set of 193 species, 122 genomes, and 111 transcriptomes. In addition, a second set of 61 transcripts from 51 hemimetabolous insect species was compiled. Accession numbers of all genome and transcriptome assemblies are listed in supplementary Table S1.

### Identification of MBD2/3 candidate sequences

Candidate methyl-CpG-binding proteins (MBDs) were identified from genome and transcriptome data using the tool tblastn (BLAST+ version 2.14.0+, (Altschul et al. 1997)) and the two described methyl-CpG-binding protein isoforms, MBD2/3-S and MBD2/3-L, from *Bombyx mori* (Xu et al. 2021; Fu et al. 2023) as queries. The resulting genomic regions covering the candidate MBD2/3 hits and 3 kb flanking sequence on both sides, were extracted from their corresponding genomes and transcriptomes. To ensure the completeness of the sequences, candidate MBD2/3 loci were manually curated. Transcripts were processed with Exonerate (version 2.4.0) (Slater and Birney 2005) to detect and translate the largest open reading frame. Protein alignments of translated genes and transcripts were computed using MUSCLE (5.1.linux64 (Edgar 2004)) and edited manually. The different isoforms of MBD2/3 were identified based on similarity to either MBD2/3-S or MBD2/3-L query and by visual examination of the protein alignment for the completeness of the MBD domain. Only a single MBD gene was identified per species examined.

### Gene Structure Prediction of Candidate MBD genes

Identification of exon-intron junctions was carried out with Exonerate (version 2.4.0) (Slater and Birney 2005) using the longer MBD2/3 isoform of a manually selected species from each order as the reference sequence of the respective order, as well as the *B. mori* reference MBD2/3-L. MBD2/3 isoforms from the following species were used as reference sequences: *B. mori* – Lepidoptera (moths and butterflies), *A. fuscipes* – Trichoptera (caddisflies), *T. castaneum* – Coleoptera (beetles), *C. cinctus* – Hymenoptera (ants, bees, wasps), *C. carnea* – Neuroptera (net-winged insects), *S. lutaria* – Megaloptera (alderflies, dobsonflies, fishflies), *I. crassicornis* – Raphidioptera (snakeflies), *D. melanogaster* and *B. oleae* - Diptera (true flies), *C. felis* - Siphonaptera (fleas), *P. vulgaris* - Mecoptera (scorpionflies), *X. vesparum* - Strepsiptera.

### Insect Phylogeny and Divergence Time

The insect phylogeny shown in Figure 2 is a compilation of phylogenetic information from the following studies. Insect topology and divergence times are based on Kjer et al. (2016) and have been extended by phylogenetic information for the orders Hymenoptera (Peters et al. 2017), Coleoptera (McKenna et al. 2019), Lepidoptera (Kawahara et al. 2019), Trichoptera (Ge et al. 2024; Frandsen et al. 2024), Diptera (Wiegmann et al. 2011) and Neuropterida (Neuroptera, Megaloptera, Raphidioptera) (Vasilikopoulos et al. 2020).

## Results

The recurrent loss of DNA methyltransferases and reduced levels or complete loss of CpG methylation, particularly in Holometabola, raises the question about the fate of MBD2/3, the sole known reader of CpG methylation in insects. To investigate this, we analyzed genomic and transcriptomic sequence data from all eleven extant holometabolous orders, using the well-described methyl-CpG-binding protein isoforms MBD2/3-S and MBD2/3-L from *Bombyx mori* as references (Fu et al. 2023; Xu et al. 2021). We used genomic and transcriptomic data to examine the exon-intron structure of the genes and transcriptome data to determine the presence of the two isoforms from a broad phylogenetic range of species.

The long isoform (MBD2/3-L) contains a complete methyl-CpG-binding domain (MBD), which is necessary for binding methylated CpGs and situated across the first two exons in *B. mori*. In contrast, the short isoform (MBD2/3-S) lacks the second exon and therefore half of this domain, a region required for structural stability and therefore mCpG binding. We will refer to this exon as exon k in the following (see Fig. 1). To place our findings in a broader evolutionary context, we also examined transcripts of representatives from all non-holometabolous insect orders as an outgroup.

**Figure 1.**
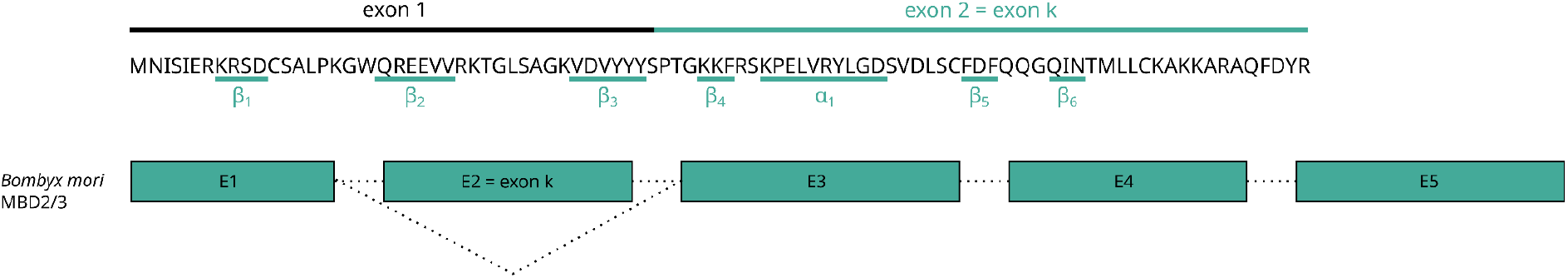
Sequence and gene structure of *Bombyx mori* MBD2/3, used as reference. Shown is the amino acid sequence encoded by the first two exons of the MBD2/3 gene, containing the MBD domain, with β-turns and α-helix highlighted (Fu et al. 2023). Below is the corresponding schematic gene structure. Exon k is alternatively spliced out to form MBD2/3-S. Alternative splicing is indicated by dotted lines below the exon.

**Figure 2.**
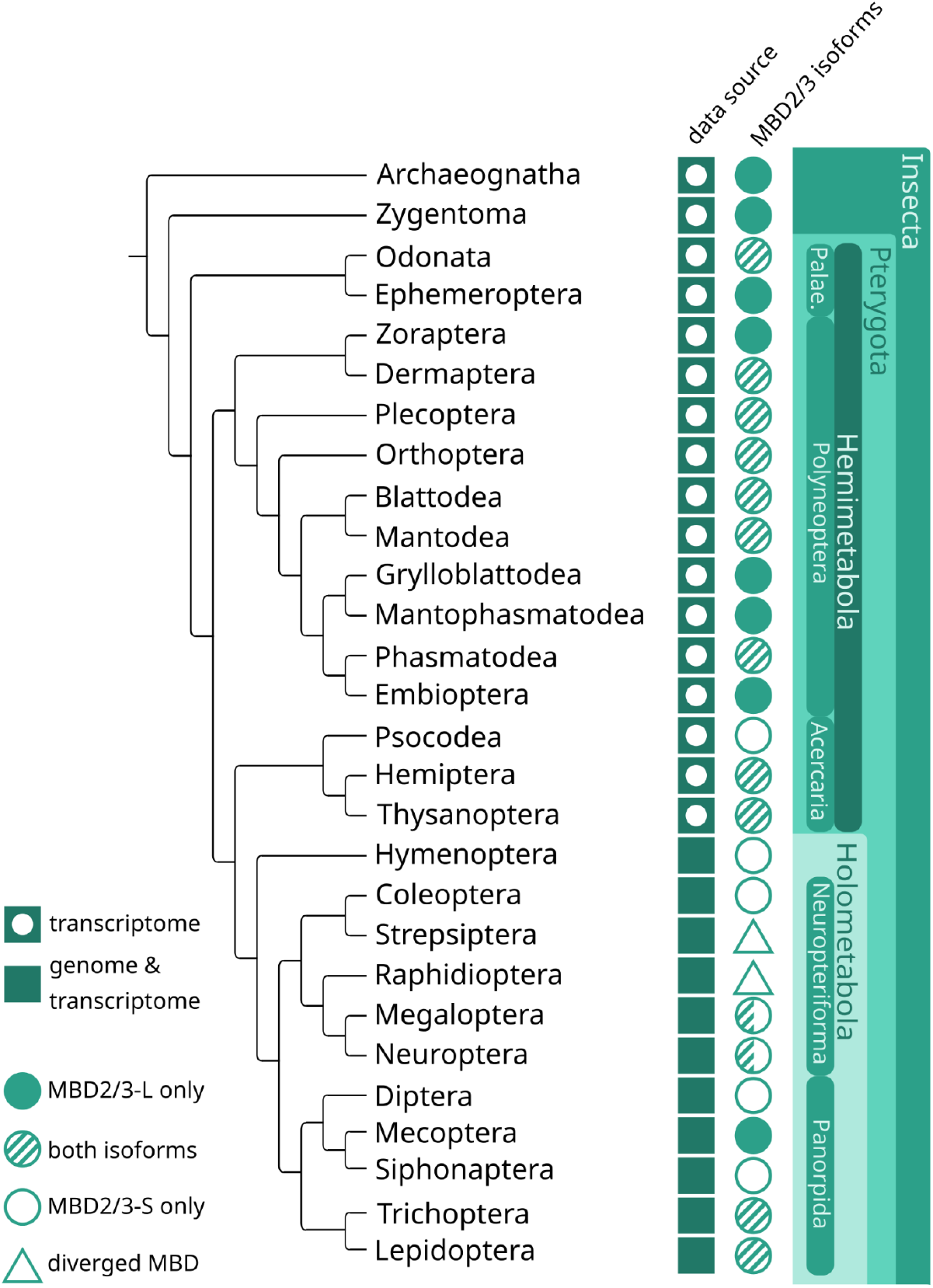
Occurrence of MBD2/3-S and MBD2/3-L isoforms across insect orders. The figure summarizes the types of data analyzed (genomic and transcriptomic) and the MBD2/3 isoforms identified across all insect orders. In some orders, the MBD domain shows divergence from the canonical form (marked with a triangle). Notably, in Megaloptera and Neuroptera, both MBD2/3-L and MBD2/3-S isoforms are present in some families, while others encode only an MBD2/3-S-like gene. Phylogenetic relationships follow the topology presented by Kjer et al. (2016).

Our analysis revealed several distinct patterns in the evolutionary fate of MBD2/3, with three major cases affecting the methyl-CpG-binding domain.

### Case 0: MBD2/3 contains a complete methyl-CpG binding domain

Within the hemimetabolous insects, we identified MBD2/3 isoforms of 51 species across all 17 extant orders. Here, we identified the long isoform MBD2/3-L in 16 of the orders, with Psocodea being the sole exception. The short isoform MBD2/3-S appears to be less prevalent, with seven orders lacking this isoform. In total, we identified 61 MBD2/3 sequences, with 41 classified as MBD2/3-L and 20 as MBD2/3-S. In Odonata (dragonflies and damselflies) and Ephemeroptera (mayflies), an insertion of 42 to 57 nucleotides following the MBD domain can be found.

We observed a similar pattern as in our reference *B. mori* in other Lepidoptera, Trichoptera, as well as in some families of Neuroptera and Megaloptera. In Lepidoptera and sister order Trichoptera, genomic analysis of 21 and 19 species, respectively, revealed that MBD2/3 is consistently encoded with a complete MBD domain. Transcriptomic data from 25 lepidopteran and 10 trichopteran species confirmed that both isoforms are commonly expressed, with few exceptions (see supplementary Table S1). The gene structure is highly conserved across all species of both orders, with perfectly conserved splice sites. As in *B. mori*, the gene consists of five exons, with the second exon alternatively spliced out to produce the short isoform MBD2/3-S (Fig. 3).

**Figure 3.**
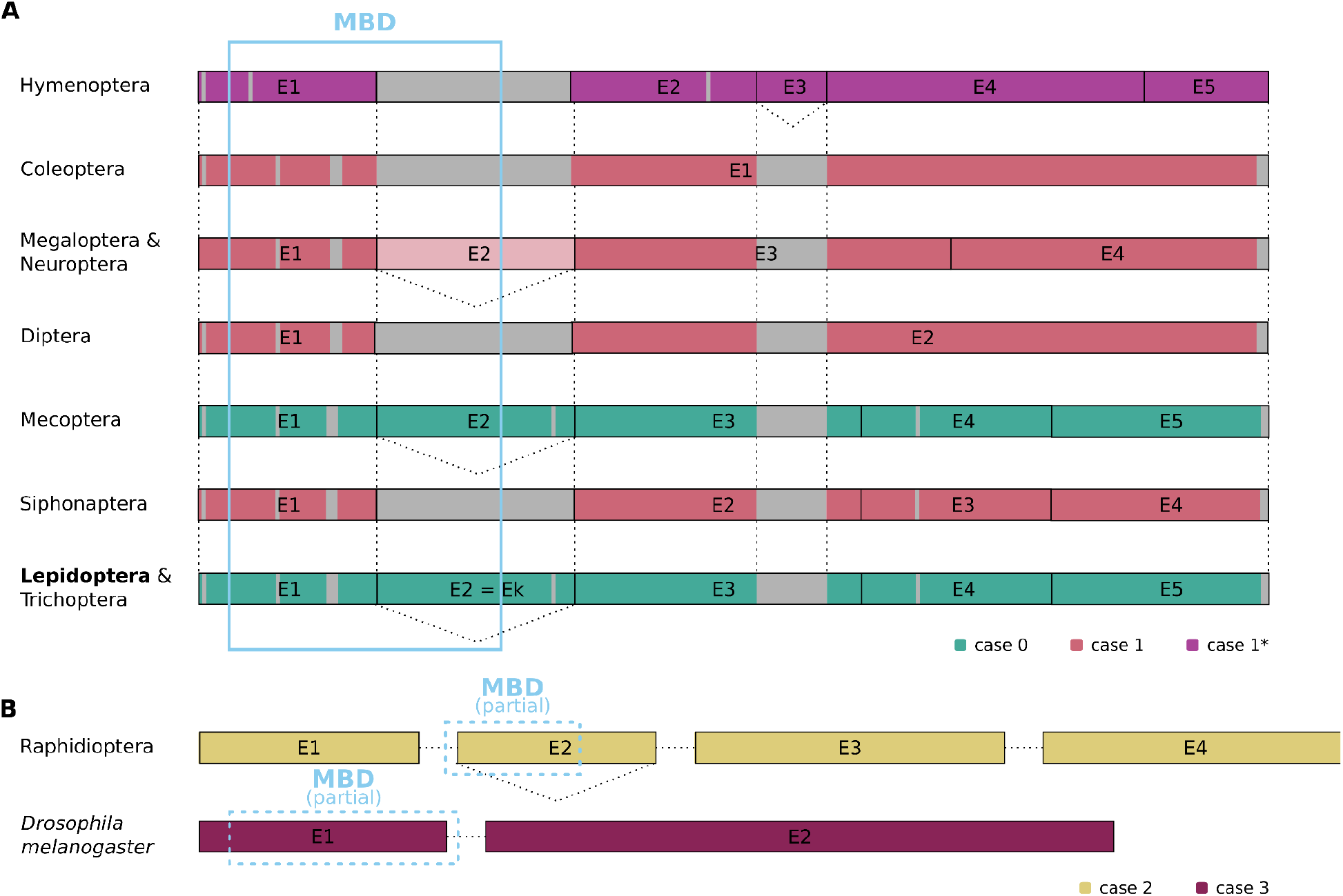
A) Alignment of the schematic representation of the MBD2/3 gene structure in Holometabola orders. Exon lengths (boxes) are shown to scale, alternative splicing is shown with dotted lines below the exons. MBD domain is highlighted with a blue box, gaps are shown in grey. Exon 2 is of lighter color in Neuroptera and Megaloptera to indicate its absence in Sisyridae and Corydalidae. Raphidioptera, *Drosophila* and Strepsiptera are not shown, as the first exon is different from all other orders and can not be aligned properly. Diptera with *Drosophila* excluded. Coleoptera: only single exon gene shown. Colors indicate the respective MBD2/3 modification cases defined in the analysis. Case 1* refers to the loss of exon k but gain of an additional exon following the MBD. **B) Schematic representation of MBD2/3 gene structure in Raphidioptera and *D. melanogaster***. Colors indicate the respective MBD2/3 modification cases defined in the analysis. Partial MBD domains are highlighted with a blue box.

This is also true for several, but not all, families of the Neuroptera and Megaloptera. We analyzed four neuropteran and three megalopteran genomes, alongside transcriptomes from 31 neuropteran and four megalopteran species. The MBD2/3 gene consists of four exons with conserved splice sites across both orders (Fig. 3). As in Lepidoptera and Trichoptera, the second exon is alternatively spliced out for the short isoform MBD2/3-S, which is frequently expressed alongside MBD2/3-L.

Similarly, in Mecoptera, we identified MBD2/3-L in the available genome data, with transcriptomic data confirming the expression of only MBD2/3-L. However, due to the limited amount of data for this order, further confirmation across additional species is needed.

### Case 1: MBD2/3 lost second half of the MBD

In the other families of Neuroptera and Megaloptera, only MBD2/3-S is encoded in the genome, lacking the second half of the MBD domain. The loss of MBD2/3-L appears to be common in these orders and confirmed in the genomes of Sisyridae (spongeflies) and in the dobsonfly species *Neoneuromus ignobilis*. In *N. ignobilis*, MBD2/3 is encoded by a single exon gene. However, in Sisyridae, the gene retains a structure similar to that of the other Neuroptera and Megaloptera in Case 0, but with exon k missing, while the splice sites remain conserved (Fig. 3). Transcriptome analysis further supports the frequent absence of the long isoform, which we did not detect in eight of the 16 families studied. No alternative transcripts were observed.

The loss of MBD2/3-L is also evident in the hemimetabolous order Psocodea (bark lice, book lice and parasitic lice) and in the holometabolous orders Siphonaptera and most Diptera. We identified only MBD2/3-S in the only available genome of Siphonaptera. This is supported by the transcriptomic data of three additional species. Most Diptera species, with the exception of *Drosophila*, also fall into this category, as we only identified MBD2/3-S, with no evidence of exon k in the genomes and transcriptomes analyzed. In four Diptera, remnants of exon k can be found, though the sequences would contain premature stop codons. In three dipteran species, a gene with a sequence similar to exon k was identified. For *Beris chalybata*, splice site predictions with the *B. mori* reference indicated two possible isoforms: a short variant matching MBD2/3-S, and a longer transcript containing a highly divergent region in place of exon k. However, no additional splice sites were predicted downstream of the predicted sequence of the MBD domain, making alternative transcripts similar to those found in other Diptera unlikely. No transcriptomes of these species are available to verify the existence of both isoforms. In the NCBI Transcriptome Shotgun Assembly database, no transcripts containing a whole MBD2/3 sequence were found.

We observed a particularly striking pattern in Coleoptera, where MBD2/3-L is absent in all species analyzed. Genomic data from 26 beetles, representing all families for which genome data were available, revealed the presence of a single MBD2/3-like gene with no paralogs. In contrast to other holometabolous orders, the MBD2/3 gene structure in Coleoptera differs between families. In 17 species, MBD2/3 is encoded by a single-exon gene (see Fig. 3), whereas in nine species, the genes contained one or two introns. The splice sites of those genes differ between families, which suggests that introns were independently acquired in several familie. In *Agrilus planipennis* and *Sitophilus oryzae*, we identified two identical copies of the MBD2/3-S gene. Given their sequence identity, we attribute this duplication to genome assembly artifacts rather than the presence of true paralogs.

Like Coleoptera, Hymenoptera only carry a gene encoding MBD2/3-S. Our genomic analysis of 16 hymenopteran species revealed a gene structure consisting of five exons but lacking exon k. Interestingly, the third exon is completely unique to Hymenoptera. Transcriptome data shows that this third exon, consisting of only 54 nucleotides, is often alternatively spliced out to form a shorter isoform, the usual MBD2/3-S. The intronic region following the first exon varies greatly in length, ranging from 93 bp up to 3664 bp. Further investigation of this intronic region revealed no remnants of exon k or conserved motifs indicative of a diverged coding sequence.

### Case 2: MBD2/3 lost first half of the MBD

We found the MBD2/3 sequence to be highly conserved within and across insect orders. Raphidioptera retained exon k, however, they have a completely different first exon, which is highly conserved within the order. The sequence is unique to Raphidioptera and does not show any similarity to the expected sequence found in other insect species. Additionally, this sequence shows no similarity to any other sequence across the tree of life in NCBI databases, suggesting a newly evolved rather than diverged exon. The first exon typically contains the first half of the methyl-CpG binding domain with β-turns β_1_-β_3_, which are essential for the formation of the MBD2/3-mCpG complex (Fu et al. 2023).

Despite this divergence, the rest of the sequence is highly similar to that of other Neuropterida, with conserved splice sites across Raphidioptera, Neuroptera, and Megaloptera. Transcriptomic data suggests that a short isoform is rarely expressed, as we only identified a short isoform in the species *Inocellia japonica*. In the remaining species, only a long isoform containing exon k was identified, making the Raphidiidae the only family within Neuropterida that lacks a short isoform.

### Case 3: MBD2/3 lost second half of the MBD and shows a diverged first half

We observed this third case exclusively in *Drosophila* and the two available Strepsiptera genomes. Notably, both Diptera and Strepsiptera have lost DNMT1 and DNMT3, leading to the complete absence of CpG methylation. In *D. melanogaster*, the sequence of the first half of the MBD domain is highly divergent compared to all other insects. The gene contains an insertion of 57 nucleotides between the β-strands β_2_ and β_3_ of the MBD domain.

In contrast, Strepsiptera lost exon k and diverged substantially in their first exon, with the first exon being significantly shorter and showing no remnants of the crucial β-strands.

### MBD2/3 lost its methyl-CpG binding ability in multiple orders of Holometabola

In a phylogenetic context, our analysis shows that while MBD2/3 remains highly conserved in sequence, it has undergone several changes throughout Holometabola, though the MBD2/3 sequence is highly conserved. In Coleoptera, Hymenoptera, Siphonaptera, and families within Neuroptera, Megaloptera, and Diptera, the long isoform (MBD2/3-L), which contains a complete methyl-CpG-binding domain (MBD), is absent. In these orders, only MBD2/3-S which lacks the second half of the MBD domain, was identified in genomes and transcriptomes.

In Raphidioptera, MBD2/3 retained a long isoform, but the first exon, which encodes the β_1_–β_3_ strands of the MBD domain, exhibits substantial sequence divergence.

In *Drosophila* and Strepsiptera, both lacking CpG methylation, the sequence of the MBD domain differed from those found in other insects. Drosophila exhibits an insertion into the β_2_ and β_3_ of the MBD domain, while the remaining Diptera show no such divergence, but a loss of exon k. In Strepsiptera, remnants of exon k are retained in the MBD2/3 gene.

Taken together, these results show that across Holometabola, MBD2/3 exhibited multiple independent modifications affecting the MBD domain. Proposed loss events of the usual MBD2/3 gene, phylogeny, divergence times and presence of DNMT1, DNMT3, and CpG methylation within the respective order are detailed in Fig. 4.

**Figure 4.**
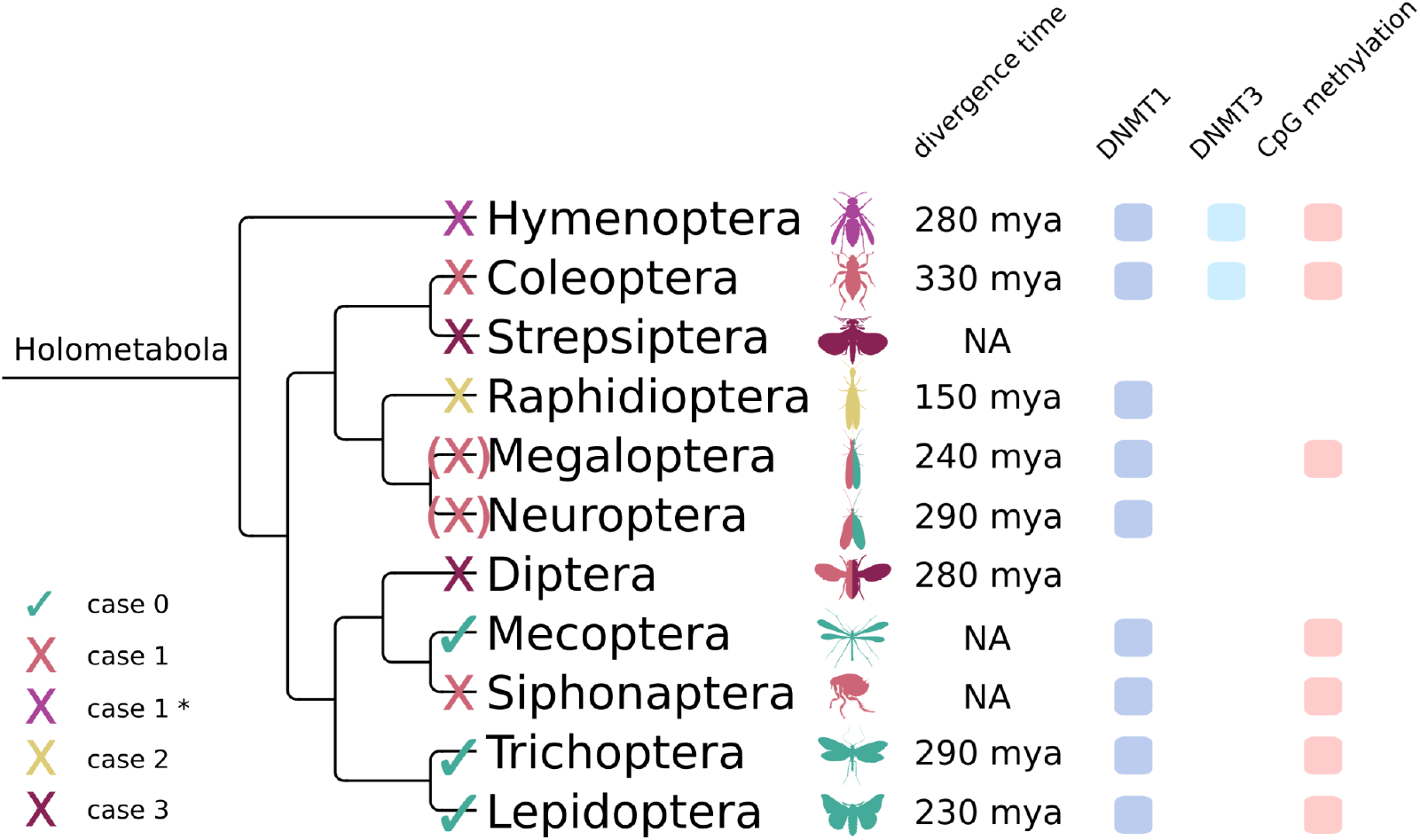
Proposed loss events of MBD2/3-L in Holometabola. The phylogeny of Holometabola is shown with evolutionary events related to MBD2/3, as proposed in this study. X marks the loss of the canonical MBD2/3-L, while a tick marks the presence. Case 1* refers to the loss of exon k but gain of an additional exon following the MBD. An (X) denotes losses observed in some, but not all, families of a given order. Colors indicate the respective MBD2/3 modification cases defined in the analysis. Phylogenetic relationships follow the topology presented by Kjer et al. (2016). Divergence times within each order are shown. Divergence times according to: Hymenoptera - Peters et al. (2017); Coleoptera - McKenna et al. (2019); Lepidoptera - Kawahara et al. (2019); Trichoptera - Ge et al. (2024); Diptera - Wiegmann et al. (2011); Neuroptera, Megaloptera, Raphidioptera - Vasilikopoulos et al. (2020). Presence of DNMT1, DNMT3, and CpG methylation according to Provataris et al. (2018), though order-level data may not capture species-specific differences, as shown by Engelhardt et al. (2022).

## Discussion

To our knowledge, this study is the phylogenetically most diverse analysis of the sequence diversification of MBD in holometabolous insects - a supraorder with a remarkably diverse CpG methylation (Provataris et al. 2018; Engelhardt et al. 2022). We found a diverse set of evolutionary modifications to the MBD2/3 gene across Holometabola. The potential loss of MBD binding ability occurred multiple times during holometabolan phylogeny independently through three different modifications to the gene. There is no consistent pattern of the presence or absence of CpG methylation coinciding with changes in the MBD2/3 domain. However, different cases of retention, modification, or loss of the methyl-CpG-binding domain in different lineages provide further insights into the diversity of the CpG methylation writer and reader toolkit in insects.

In invertebrate genomes, the ancestral MBD2/3 protein is encoded by a single gene, in contrast to the presence of MBD2 and MBD3 in vertebrates, which arose following a duplication event (Hendrich and Tweedie 2003). Previously, it has been reported that insects can possess two structurally and functionally distinct isoforms, MBD2/3-L and MBD2/3-S. Contrary to MBD2/3-L, the short isoform is unable to bind methylated CpG sites through a loss of an exon encoding structurally relevant parts of the MBD domain, specifically the β_4_–β_6_ strands and α_1_ helix (Fu et al. 2023). Those structures critically affect the stability of the actual MBD-DNA binding site encoded on exon 1 (Fu et al. 2023). The absence of this second half of the MBD domain, encoded in what we call exon k, has been shown to disrupt the CpG binding ability in *B. mori* (Lepidoptera) and *T. castaneum* (Coleoptera) (Song et al. 2020; Xu et al. 2021). Most hemimetabolous and some holometabolous orders, such as Lepidoptera, still retain both isoforms. Functional differentiation of those isoforms was shown in *Bombyx mori*, where only the long isoform is critical for embryonic development (Fu et al. 2023).

In contrast, we found that Coleoptera, Hymenoptera, parts of Neuroptera and Megaloptera, Siphonaptera, and the hemimetabolous Psocodea encode only the short MBD2/3 isoform, lacking exon k and thus mCpG binding ability. In Coleoptera, we propose the origin of the single-exon gene encoding only MBD2/3-S through retrotransposition in the germ line in an early common ancestor of Coleoptera. The pseudogene then gained potential beneficial regulatory elements (Wicker et al. 2007) and ultimately replaced the original MBD2/3, which was spliced to both isoforms and likely accumulated mutations until functional loss. Additional introns found in some families were later independently acquired. Despite the loss of MBD2/3-L, the expression patterns in *T. castaneum* show similarities to those o*f B. mori* MBD2/3 (Fu et al. 2023), with the highest expression in early development (Song et al. 2020), implying no drastic change in its methylation-independent function.

Conversely, certain losses in CpG methylation or the ability to read those marks might be disadvantageous. While Hymenoptera lost exon k in the early evolution of this order, the function of MBD2/3 may have been restored through the emergence of a third exon, whose role remains unclear but may stabilize the protein structure similarly to exon k, thereby potentially restoring or altering the binding affinity. There are contrasting reports on the MBD2/3 mCpG binding ability in *Apis mellifera*. While Wang et al. (2006) report mCpG binding ability of the longer splice variant, Liu et al. (2019) did not confirm this. These changes might reflect functional compensation, add a new function, or they may represent neutral events with no selective advantage. However, given the high sequence conservation within Hymenoptera, even with an estimated divergence time of 280 mya (Peters et al. 2017) in the selected species, we hypothesize that this additional exon has an important function, either for restoring mCpG binding ability or other neofunctionalizations.

Contrary, Raphidioptera lost exon 1, and thereby the sequence encoding the DNA-MBD2/3 interaction site, entirely and acquired a novel first exon with no homology to known insect sequences. Given the absence of CpG methylation in this order (Provataris et al. 2018), this raises the question whether the unique structure of exon 1 is an adaptation towards a novel function of MBD2/3 in the absence of DNA methylation (binding of mCpT/A instead of mCpG), as it could be the case in the dipteran genus *Drosophila* (Marhold et al. 2004).

Notably, no MBD2/3-L-like transcripts have been identified in any Dipteran species. In eight species, remnants of exon k were detected with varying levels of sequence divergence. Those remnants seem to either be rendered intronic or, in four species, are predicted with inconsistent splice sites. Splice variants of those genes would contain remnants of the sequence that might impact mCpG binding ability. However, in the alternative prediction of an MBD2/3-S transcript, the conserved splice sites associated with MBD2/3-S in Diptera were consistently recovered, making these predictions the more plausible result. Due to the absence of transcriptomic data for these species, we cannot confirm the existence or functional relevance of these predicted variants at this point. Therefore, we conclude that exon k diverged in Diptera until its eventual loss of function.

Mecoptera retain the full MBD2/3-L gene, while Siphonaptera have lost exon k. Strepsiptera encode an MBD2/3 gene missing exon k and showing a diverged sequence in the first half of the MBD domain. It is unclear whether the loss of exon k preceded the divergence of Strepsiptera and Coleoptera, or if the loss occurred independently in those orders. However, data from these groups remain limited, and more complete genomes and transcriptomes will be necessary to draw firm conclusions.

While the MBD domain in several orders of Holometabola diverged until the eventual loss of mCpG binding ability, other parts of MBD2/3 remain highly conserved. Besides the intrinsically disordered region (IDR), which can mediate protein-protein interactions (Roterman, Stapor, and Konieczny 2023; Tompa et al. 2015), the coiled-coil domain, essential for NuRD complex formation (Leighton and Williams 2020), remains conserved as well. This MBD2/3 domain connects the two functional distinct NuRD subcomplexes, specifically through binding the HDCC and the GATAD2 protein, an interaction site that has emerged with the earliest multicellular organisms (Cramer et al. 2017). Among other functions, the chromatin-remodeling NuRD complex regulates transcription, is involved in DNA repair, and is equally important in methylation dependent and independent contexts (Ramírez et al. 2012; Leighton and Williams 2020; Smeenk et al. 2010; Smeenk and van Attikum 2011; Z. Liu et al. 2024; O’Shaughnessy and Hendrich 2013). Notably, the MBD domain itself is not considered necessary to form the NuRD complex (Leighton and Williams 2020).

In most vertebrates except Xenopus (Wade et al. 1999), MBD3 lost its CpG binding ability due to the point mutations of two key amino acids within the MBD domain (Du et al. 2015), but retains NuRD associated roles. Early findings linked MBD3 to lineage commitment, while later studies showed conflicting roles in promoting or inhibiting pluripotency and reprogramming, likely due to an incomplete molecular understanding of NuRD function (Leighton and Williams 2020). In mammalian adults, MBD3 mediated methylation-independent NuRD functions are likely prominent in undifferentiated tissues, whereas methylation-dependent roles are critical during differentiation. However, the lack of mechanistic insight into NuRD-mediated gene regulation limits interpretation of these patterns (Leighton and Williams 2020).

Despite the inherent limitations of comparing vertebrate MBD2 and MBD3 to invertebrate MBD2/3, with and without complete MBD domains, vertebrate studies may offer valuable insights. In vertebrates, the NuRD-MBD2 complex is linked to gene repression, while the NuRD-MBD3 complex is associated with active genes (Shimbo et al. 2013). CpG methylation in invertebrates is also associated with active gene expression (Lewis et al. 2020). These differences could suggest that methylation-independent NuRD functions may be more relevant in insect lineages with reduced CpG methylation, as it is the case in holometabolous insects. Furthermore, in holometabolous insects, reduced CpG methylation levels may have favored the loss of the MBD mCpG binding ability, as maintaining a complex for reading sparse and potentially noisy mCpG methylation signals might be maladaptive.

A variable pattern of DNMT loss and CpG methylation reduction has been observed in Coleoptera (Engelhardt et al. 2022). Interestingly, we detect the loss of mCpG binding ability in the CpG methylation reader at the very origin of Coleoptera. Potentially, the loss of this reader, coupled with an overall low CpG methylation level resulted in an increasing redundancy of methylation, contributing to the loss in certain species, while others preserved CpG methylation. A markedly different pattern emerges in Neuroptera, which are predicted to lack CpG methylation altogether (Provataris et al. 2018), yet some families have retained MBD2/3-L. These contrasting patterns highlight the diverse and lineage-specific ways in which components of the CpG methylation machinery have been lost, underscoring that there is no single evolutionary path shared across insect orders, but all may reflect shifts away from a CpG methylation based gene regulation.

## Conclusions

Our findings reveal a diverse set of evolutionary modifications to the MBD2/3 gene across insect orders, particularly in Holometabola, where CpG methylation is reduced or lost entirely. The retention, modification, or loss of the methyl-CpG-binding domain of the MBD2/3 protein in different lineages provides further insights into the diversification of the Insect CpG methylation toolkit and emphasizes the importance of avoiding broad generalizations across orders.

## Acknowledgments

This study was funded by the Deutsche Forschungsgemeinschaft (DFG, German Research Foundation) – priority program SPP 2349, project number 503 349 225 to S.J.P. and J.K.

## Declaration of Interests

The authors declare no conflict of interests.

## Notes

### Competing Interest Statement

The authors have declared no competing interest.

